# Stability, affinity and chromatic variants of the glutamate sensor iGluSnFR

**DOI:** 10.1101/235176

**Authors:** Jonathan S. Marvin, Benjamin Scholl, Daniel E. Wilson, Kaspar Podgorski, Abbas Kazemipour, Johannes Alexander Müeller, Susanne Schoch-McGovern, Francisco José Urra Quiroz, Nelson Rebola, Huan Bao, Justin P. Little, Ariana N. Tkachuk, Adam W. Hantman, Samuel S.-H. Wang, Edwin R. Chapman, Dirk Dietrich, David A. DiGregorio, David Fitzpatrick, Loren L. Looger

## Abstract

Single-wavelength fluorescent reporters allow visualization of specific neurotransmitters with high spatial and temporal resolution. We report variants of the glutamate sensor iGluSnFR that are functionally brighter; can detect sub-micromolar to millimolar concentrations of glutamate; and have blue, green or yellow emission profiles. These variants allow *in vivo* imaging where original-iGluSnFR was too dim, reveal glutamate transients at individual spine heads, and permit kilohertz imaging with inexpensive, powerful fiber lasers.

## Main Text

The intensity-based glutamate-sensing fluorescent reporter (iGluSnFR)^1^ has become an invaluable tool for studying glutamate dynamics in diverse systems, including retina^2,3^, olfactory bulb^4^ and visual cortex^5^. iGluSnFR also allows mesoscale “functional connectomic” mapping^6^ and mechanistic studies of Huntington’s disease^7^, synaptic spillover^8^, cortical spreading depression^9^ and exocytotic vesicle fusion^10^. However, iGluSnFR is insufficient for some applications due to poor expression (in some preparations), and kinetics that do not match the time courses of some phenomena. Here we describe variants that are functionally brighter (due to increased membrane expression), have tighter or weaker affinity, and fluoresce blue, green, or yellow.

Replacement of circularly permuted eGFP with circularly permuted “superfolder” GFP^11^ (SF-iGluSnFR) yielded 5-fold higher soluble-protein expression levels in bacteria (0.5 μmol/1L growth *vs.* 0.1 μmol/1L). Circular dichroism indicates an increase in melting temperature transition (T_m_) of∼5°C (**Supp. Fig. S1**). The 2-photon cross-section and excitation, emission, and absorption spectra of SF-iGluSnFR are similar to the original (**Supp. Fig. S2a-d**). Head-to-head comparison of SF-iGluSnFR with original-iGluSnFR in mouse somatosensory cortex shows substantially more robust expression by the former (**Supp. Fig. S3a,b**). Under typical imaging conditions (<20 mW, 130-nanosecond dwell time per pixel), SF-iGluSnFR is bright enough for repeated imaging, while original-iGluSnFR is too dim (**Supp. Fig. S3c,d**). While we observed a faster 2-photon *in vivo* photobleaching rate for SF-iGluSnFR in somatosensory cortex (**Supp. Fig. S3e**), partially-bleached SF-iGluSnFR was still brighter than iGluSnFR. Thus, SF-iGluSnFR will have superior expression *in vivo,* where the quantity of deliverable DNA can be limiting.

While the affinity of membrane-displayed iGluSnFR (4 μM) is adequate for some *in vivo* applications, tighter variants are needed for circumstances of limiting glutamate concentrations, *e.g.* at sparsely-firing synapses. Additionally, measuring glutamate release events with raster scanning requires variants with slower off-rates so that the decay time from glutamate binding is long enough to be sufficiently sampled at the operating frame rate (typically <100 Hz). Replacement of eGFP with superfolder GFP increases the *in vitro* affinity of soluble SF-iGluSnFR over original-iGluSnFR (40 μM vs. 80 μM, **Supp. Fig. S4a**). To further modulate affinity, we exploited the conformational coupling between the open-closed equilibrium of bacterial periplasmic binding proteins (PBPs, *e.g.* the glutamate-binding protein in iGluSnFR) and their ligand-binding affinity^12^. Briefly, mutation of residues in the “hinge” of PBPs can allosterically alter affinity, without compromising the stereochemical integrity of the ligand-binding site. In a bacterial lysate assay, we screened an A184X library of the iGluSnFR glutamate-binding domain (mutated to valine in the original-iGluSnFR). Reversion to alanine or other small amino acids tightened affinity, while larger side chains weakened affinity (**Supp. Fig. S5**).

We introduced A184S into SF-iGluSnFR to generate a tighter variant. (Reversion A184A had low ΔF/F.) Affinities of purified soluble protein are 7 μM and 40 μM for the A184S and A184V (unmutated from iGluSnFR) SF-iGluSnFR variants, respectively (**Supp. Fig. S4a**). The tighter affinity of A184S arises from a slower off-rate (**Supp. Fig. S4b**). The affinity variants were recloned into an AAV vector containing an IgG secretion signal and a PDGFR transmembrane domain. Viral expression on cultured rat hippocampal neurons (AAV2/1.*hSynapsin1*. SF-iGluSnFR) yields glutamate affinities∼10x tighter than the soluble form (0.7 μM and 2 μM for A184S and A184V, respectively; **Supp. Fig. S6**). A similar increase in affinity upon membrane tethering was seen with the original-iGluSnFR^1^. Whole-field stimulation (50 Hz) of these cultures shows that their relative half-times of fluorescence decay parallel their *in vitro* kinetics, with all variants having faster decay than GCaMP6f (**Supp. Fig. S7**).

*In vivo*, the tighter/slower SF-iGluSnFR.A184S variant shows improved detection of stimulus-evoked glutamate release in ferret visual cortex in response to presented drifting gratings (**Fig. 1a,b**). Peak amplitudes reached 30% ΔF/F for SF-iGluSnFR.A184S but only 5% ΔF/F for SF-iGluSnFR.A184V when imaged at 30 Hz. Greater ΔF/F of SF-iGluSnFR.A184S allows extraction of robust orientation tuning curves compared to SF-iGluSnFR.A184V. Enhanced sensitivity of A184S also allowed orientation-selective responses to be resolved in individual dendritic spines (**Fig. 1c,d**). Synaptic glutamate release as measured with SF-iGluSnFR.A184S was not only strongly selective for visual stimuli, but response amplitudes across individual trials consistently exceeded A184V over all stimulus-evoked responses (A184S median ΔF/F = 16%, n = 72 spines; A184V median ΔF/F = 9%, n = 22 spines; p = 2e-115, Wilcoxon rank-sum test) or only preferred stimuli (A184S median ΔF/F = 27%, n = 72 spines; A184V median ΔF/F = 14%, n = 22 spines; p = 9e-23) (**Supp. Fig. S8**).

**Fig. 1.**
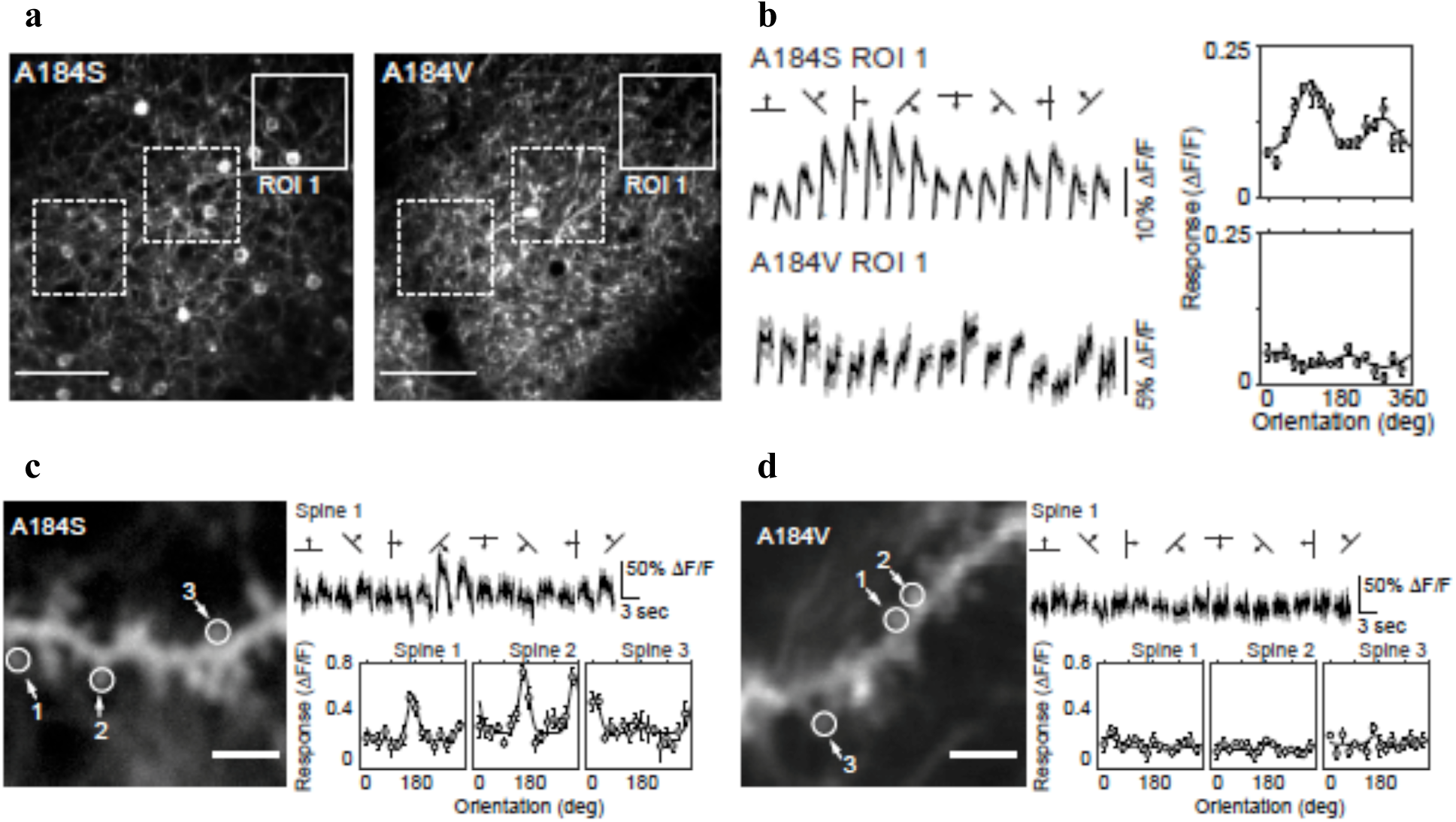
SF-iGluSnFR.A184S shows larger responses to visual stimuli than SF-iGluSnFR.A184V. a) Two-photon standard-deviation projection of SF-iGluSnFR.A184S and A184V expressed in ferret visual cortex (A184S: 190 μm, A184V: 175 μm, scale bar 100 μm). b) Trial-averaged stimulus-evoked responses (shown for ROI 1) reveal robust orientation tuning and peak amplitudes of∼30% ΔF/F for A184S. Peak responses plotted as a function of stimulus orientation show robust selectivity with the A184S variant. For the A184V variant, stimulus-evoked fluctuations are too small (∼5% ΔF/F) to generate robust tuning plots. c) Two-photon standard-deviation projection of an isolated dendritic segment with active spines revealed with SF-iGluSnFR.A184S. Individual dendritic spines are driven selectively and strongly by drifting gratings. Orientation tuning from peak responses shows large spine responses (30-50% ΔF/F) and, importantly, reveals that spines on a single dendritic branch can receive differently tuned excitatory input. d) Same as in (c) for SF-iGluSnFR.A184V. Dendritic spine responses with A184V are weak and almost unresolvable.

While slow off-rate variants of SF-iGluSnFR are better for detecting individual synaptic events by temporal summation of fluorescence, faster off-rate variants are needed for temporal resolution spiking dynamics and at large synapses where glutamate clearance is limiting. Soluble SF-iGluSnFR.S72A has 200 μM affinity for glutamate (**Supp. Fig. S4a**), arising from a combination of both slower on-rate and faster off-rate (**Supp. Fig. S4b**). In neuronal culture, S72A has an affinity of 35 μM,∼10x weaker than its parent, A184V (**Supp. Fig S6).**

In neuronal culture, fluorescence of the culture (not localized to specific structures) returns to baseline within 100 msec. of a single electrical stimulation for S72A, faster than A184V, A184S, or GCaMP6f (**Supp. Fig S7**). The substantially faster off-rate of S72A provides enhanced temporal resolution of paired (20 Hz) electrical stimuli over A184V (**Fig. 2a,b; Supp. Fig. S9**), making it useful for assessing short-term synaptic plasticity. A train of 6 electrical pulses (20 Hz) in 1 mM extracellular Ca^2+^ were resolved as equal, individual release events with S72A, while A184V yielded an integrated signal (**Fig. 2c,d**). In 3.5 mM extracellular Ca^2+^, vesicles are released with higher probability during the initial stimulation^13^. This was observed by S72A, as reported by a reduction in fluorescence response as the pulse train progresses (**Fig. 2c**), while these differences are obscured by the slower decay of A184V (**Fig. 2d**). Thus, while S72A has a lower ΔF/F in response to the same amount of glutamate being released (due to weaker affinity), its faster kinetics provides enhanced temporal resolution of synaptic activity. Similarly, S72A provides enhanced spatial resolution of glutamate release over A184V (**Supp. Fig. S10**).

**Fig. 2.**
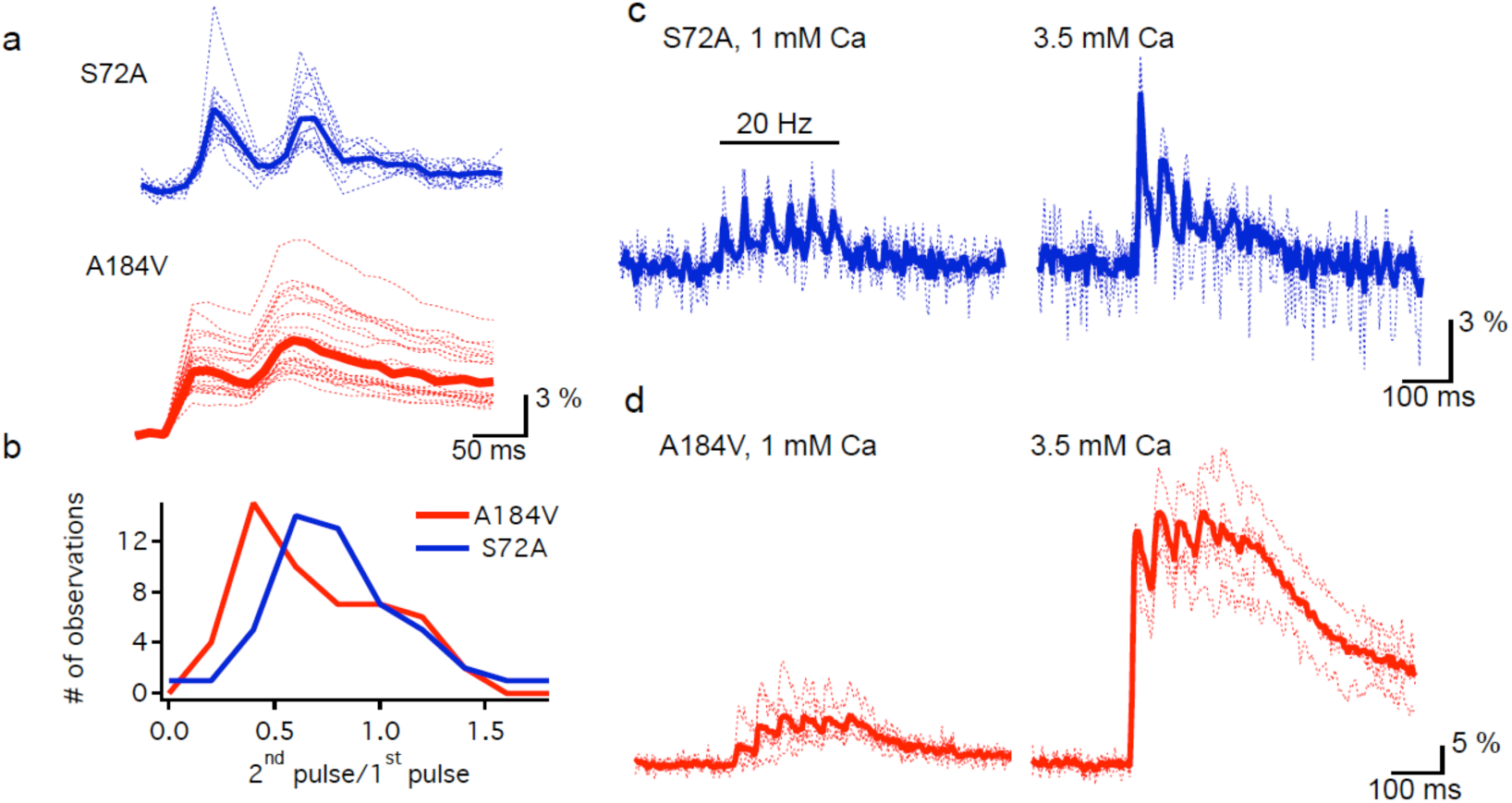
SF-iGluSnFR.S72A permits resolution of multiple glutamate release events in cultured mouse embryonic hippocampal neurons. a) Single (dashed) and averaged (solid) traces of SF-iGluSnFR.S72A (blue) and SF-iGluSnFR.A184V (red) response to 20 Hz paired electrical stimuli. b) Histogram showing intensity ratio of second pulse to first pulse. c) The faster off-rate of S72A can be used to observe short-term synaptic plasticity, potentially vesicle release depression. Higher concentrations of extracellular calcium can increase vesicle release, leading to vesicle exhaustion as the train of field pulses progresses. d) The slow decay of A184V obscures this depression.

With fast rise and decay times, we examined whether SF-iGluSnFR can serve as an alternative to GCaMP6f for monitoring neuronal activity in mouse cerebellar brain slice. Single cerebellar granule cell bouton responses to single action potentials (APs) could indeed be resolved using fast linescan detection (< 1ms per line; **Fig. 3a**), and were much faster than GCaMP6f rise and decay times at both 2 mM and 1.5 mM extracellular calcium. The S72A variant had by far the fastest response (S72 half decay 7.9 ± 1.0 ms, A184V 28.1 ± 1.6 ms, GCaMP6f 1.5 mM [Ca^2+^]_e_ 37.9 ± 3.9 ms, GCaMP6f 2.0 mM [Ca^2+^]_e_ 108.6 ± 8.8 ms). The signal-to-noise-ratios (SNRs) were best for A184V, but even S72A produced better SNRs than GCaMP6f under physiological extracellular calcium concentrations (1.5 mM). The superior SNR of A184V showed putative single vesicle release events in single trials (**Fig. 3b**). If many bouton responses are pooled and averaged for each trial, single spike detection at 20 Hz is feasible (average trace, **Fig. 3b**). For 20 Hz stimuli, both A184V and S72A produced little accumulation of bouton fluorescence after 10 stimuli as compared to GCaMP6f (**Fig. 3c**), similar to dendritic responses in culture (**Fig. 2**). At 100 Hz, discrete release events could be detected, in contrast to GCaMP6f (**Fig. 3d**). Note the poor temporal precision of the train response, in contrast to A184V and S72A. Thus, both A184V and S72A enable a larger dynamic range of reported firing frequencies, with S72A providing the largest range due to its low affinity. Moreover, the fast kinetics of SF-iGluSnFR.A184V and SF-iGluSnFR.S72A can more reliably estimate spike times (versus GCaMP6f), and are much better suited to high-frequency spike detection (> 100 Hz), as in cerebellar granule cells^14^.

**Fig. 3.**
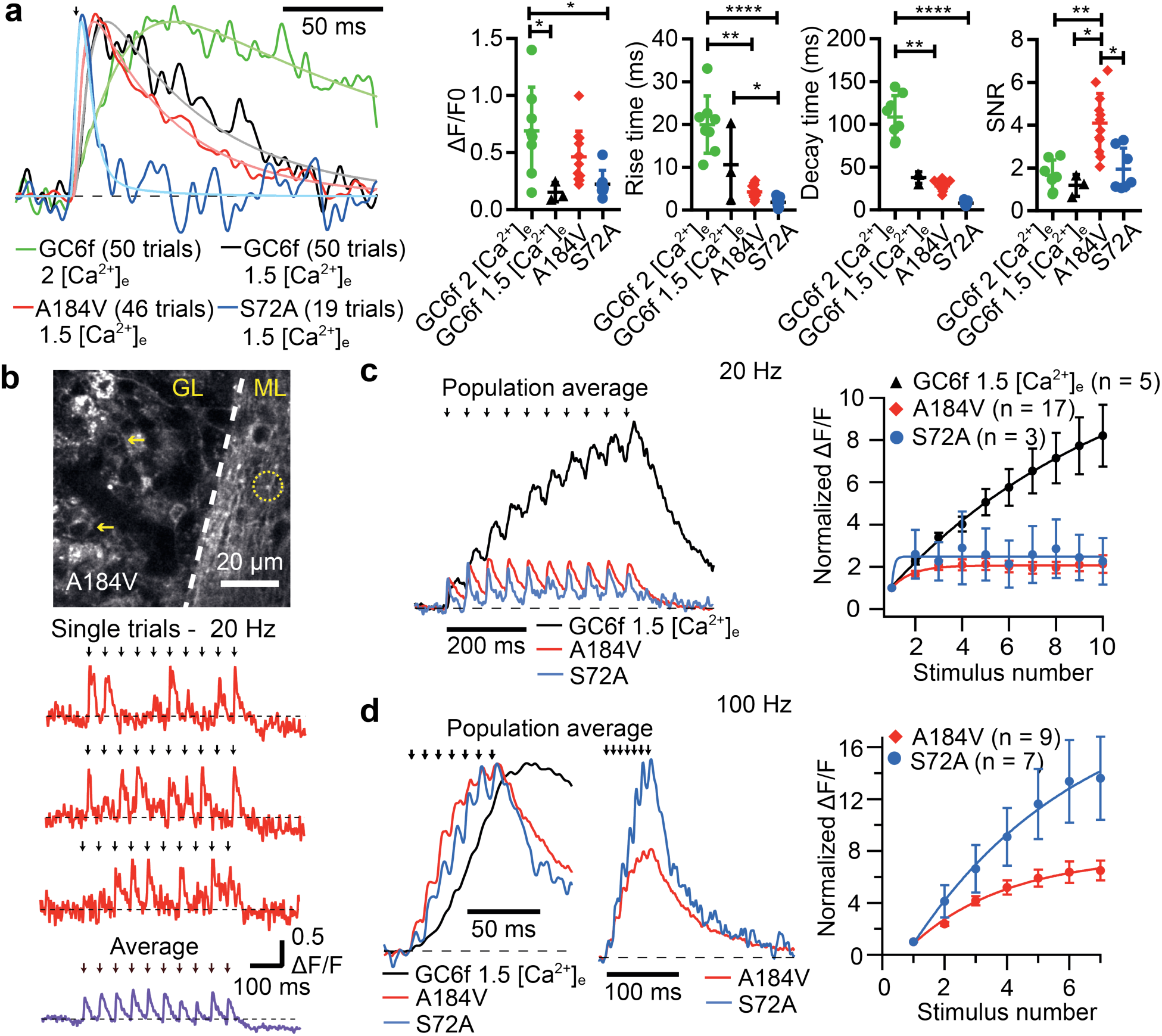
S72A variant shows faster bouton fluorescence signals resulting from single or trains of electrical stimuli in mouse cerebellar brain slice. **a)** Averaged response from single boutons expressing GCaMP6f (GC6f) at 2 mM [Ca^2+^]_extracellular_ (green), GC6f at 1.5 mM [Ca^2+^]_e_ (black), SF-iGluSnFR.A184V at 1.5 mM [Ca^2+^]e (A184V, red) and SF-iGluSnFR.S72A at 1.5 mM [Ca^2+^]_e_ (S72A, blue), normalized to peak response. In parenthesis, the number of trials used to calculate the average. Right, summary plots of ΔF/F_0_, 10-90% rise time, 50% decay time and signal-to-noise-ratio (SNR). Multiple comparisons were performed with the Kruskal-Wallis test and the Dunn’s multiple comparisons test.* *P* ̼ 0.05, ** P < 0.01, **** *P* < 0.0001. **b)** Two-photon fluorescence image of granule cells and parallel fibers expressing A184V in cerebellar slice (GL –granule layer, ML – molecular layer). Yellow arrows indicate labeled somata of granule cells, and circle indicates boutons from parallel fibers. Bottom, example of single trial A184V fluorescence responses to 20 Hz electrical stimulation (red) and the average of 10 trials (purple). **c)** Population-averaged fluorescence responses to 20 Hz stimulation (n boutons =5, GC6f; n=17, A184V; n=3, S72A). Traces are normalized to the peak of the first response. **d)** Population average of response to 100 Hz electrical stimulation (n boutons =9, GC6f; n=9, A184V; n=7, S72A) normalized to the maximum amplitude (left) or to the peak of the first response (middle), and average response of all the boutons. n is number of boutons. Black arrows indicate time of electrical stimulation.

Introduction of chromophore mutations from GFP variants Azurite^15^ or Venus^16^to SF-iGluSnFR led to functional blue and yellow versions, respectively. The former required re-optimization of residues linking the FP with the glutamate-binding protein. The latter was a straightforward modular replacement. (Annotated amino acid sequences are given in **Supp. Fig. S11**). SF-Azurite-iGluSnFR has significantly lower ΔF/F (**Supp. Fig. S12**). SF-Venus-iGluSnFR has similar affinity and maximum fluorescence response to glutamate as SF-iGluSnFR, but with red-shifted excitation and emission spectra (**Supp. Fig. S13**). Importantly, its 2-photon excitation spectrum is sufficiently red-shifted to allow strong excitation at 1030 nm (**Supp. Fig. S13**), compatible with relatively inexpensive, powerful femtosecond fiber lasers^17^. These powerful lasers enable simultaneous excitation of many foci, enabling very fast (1016 Hz) large-area imaging by recording projections of the sample and computationally reconstructing images (Kazemipour, et al., accepted). In neuronal culture, two near-simultaneous pulses of glutamate uncaging were resolved with both high spatial and temporal resolution by measuring fluorescence changes in a neuron expressing SF-Venus-iGluSnFR.A184V (**Fig. 4** and **Supp. Movie 1**).

**Fig. 4.**
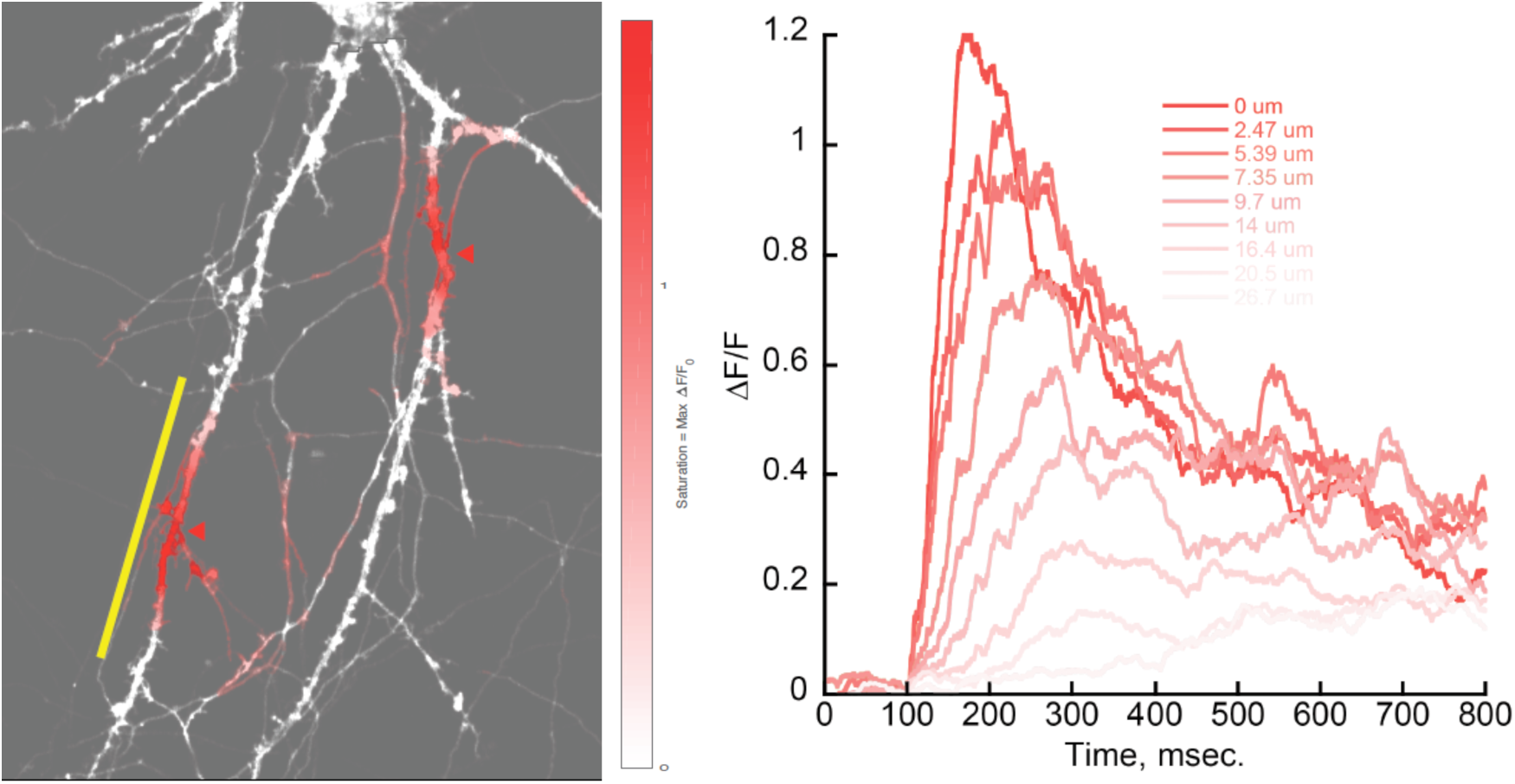
High-speed two-photon imaging (1016 Hz frame rate) of a cultured neuron expressing SF-Venus-iGluSnFR using 1030 nm excitation. Left) RuBi-glutamate was uncaged for 10 msec at each of two 5 μm spots (red arrowheads) on two adjacent dendrites. Color saturation denotes the glutamate transient amplitude (see scale bar). Yellow line indicates locations for traces shown at (right). Right) Recorded traces at nine pixels at various distances from the uncaging focus, parallel to the yellow line at (left). The traces are approximate maximum-likelihood solutions recovered with the FADE algorithm (Kazemipour et al., accepted). This recording is of a single uncaging event, without averaging. A subregion of the field of view is shown at (left). Supplemental Movie 1 shows the entire recording.

The iGluSnFR variants described here increase the power of genetically encoded glutamate imaging. Affinity variants with altered kinetics broaden the range of observable glutamate release events. Chromatic mutants allow fast imaging with cheap lasers, and potential utility in multi-color imaging. Improved membrane targeting and photostability will be valuable in all applications.

Note: Supplementary information is available in the online version of the paper.

## Accession codes

All constructs have been deposited at Addgene (#’s pending). All sequences have been deposited in Genbank (#’s pending). AAV virus will be available from Addgene and from University of Pennsylvania Vector Core (http://www.med.upenn.edu/gtp/vectorcore/).

## Acknowledgements

We would like to thank John Macklin (Janelia Research Campus) for 2-photon spectra, and Kim Ritola and Janelia Virus Services for AAV production.

## Author contributions

JSM & LLL: Protein engineering and manuscript; BS, DEW, and DF: Ferret visual cortex; JAM, SS-M, and DD: Neuronal culture analysis; JPL: Brightness assessment in somatosensory cortex; KP & AK: High speed imaging of yellow variant; ANT: Protein engineering; HB & ERC: Stopped-flow kinetics; NR, FJUQ, SW, AWH, DAD: Cerebellum.

## Reagent availability

All analysis code used in this study is available upon request.

## Code availability

All analysis code used in this study is available upon request.

## Data availability

All data from this study is available upon request.

